# Brief Communication: Magnetic Immuno-Detection of SARS-CoV-2 specific Antibodies

**DOI:** 10.1101/2020.06.02.131102

**Authors:** Jan Pietschmann, Nadja Vöpel, Holger Spiegel, Hans-Joachim Krause, Florian Schröper

**Affiliations:** Fraunhofer Institute for Molecular Biology and Applied Ecology IME, Forckenbeckstraße 6, 52074 Aachen; Institute of Biological Information Processing, Bioelectronics IBI-3, Forschungszentrum Jülich, 52428 Jülich, Germany

**Keywords:** COVID-19, frequency mixing technology, immunofiltration, magnetic beads, point-of-care diagnostic

## Abstract

SARS-CoV-2 causes ongoing infections worldwide, and identifying people with immunity is becoming increasingly important. Available point-of-care diagnostic systems as lateral flow assays have high potential for fast and easy on-site antibody testing but are lacking specificity, sensitivity or possibility for quantitative measurements. Here, a new point-of-care approach for SARS-CoV-2 specific antibody detection in human serum based on magnetic immuno-detection is described and compared to standard ELISA. For magnetic immuno-detection, immunofiltration columns were coated with a SARS-CoV-2 spike protein peptide. SARS-CoV-2 peptide reactive antibodies, spiked at different concentrations into PBS and human serum, were rinsed through immunofiltration columns. Specific antibodies were retained within the IFC and labelled with an isotype specific biotinylated antibody. Streptavidin-functionalized magnetic nanoparticles were applied to label the secondary antibodies. Enriched magnetic nanoparticles were then detected by means of frequency magnetic mixing detection technology, using a portable magnetic read-out device. Measuring signals corresponded to the amount of SARS-CoV-2 specific antibodies in the sample. Our preliminary magnetic immuno-detection setup resulted in a higher sensitivity and broader detection range and was four times faster than ELISA. Further optimizations could reduce assay times to that of a typical lateral flow assay, enabling a fast and easy approach, well suited for point-of-care measurements without expensive lab equipment.

## 1. Introduction

The new Severe Acute Respiratory Syndrome Coronavirus-2 (SARS-CoV-2), is causing ongoing worldwide infections, leading to an unprecedented pandemic. According to World Health Organization (WHO), it is estimated that up to 82% of people with coronavirus disease 19 (COVID-19) are not aware that they are/were infected due to no or very mild symptoms [1]. COVID-19 symptoms can be noted comparable to common cold cough, rhinitis or fever up to harsh symptoms, especially at elderly, such as respiratory problems with lung failure and death [2, 3]. Patients with few or no symptoms in particular pose the greatest risk, as they can infect many more people via a droplet infection [4]. Identifying people who were infected and have obtained immunity against SARS-CoV-2 is becoming highly important. Besides valuable epidemiological information regarding distribution and spreading behavior, detection methods would help to manage the currently imposed restrictions and non-pharmaceutical measures. All people with confirmed immunity most probably would have no risk to infect themselves and thus would not represent a risk of infection for others. At a later point, serological assays would also be required to prove and monitor effectivity of vaccination and longevity of the obtained immunity. Fast, cheap and easily applicable on-site testing solutions will thus become increasingly important, but currently only few rapid test systems are available. Lateral-flow-detection (LFD) approaches are easy to handle and results are gained after 10-15 min. However they are not quantitative, and their reliability, specificity and sensitivity is much worse than that of lab-based assay formats based on enzyme-linked immunosorbent assay (ELISA). In particular, specificity is a major challenge at currently available serological antibody tests. This depends to a large extent on the antigen used in the test assays. Enveloped positive-stranded RNA SARS-CoV-2 coronaviruses consist of five structural proteins, the spike glycoprotein (S), envelope protein (E), membrane protein (M), the nucleocapsid protein (N) and a hemagglutinin esterase (HE). The S-protein, a complex folded glycoprotein comprising two regions, S1 and S2, exhibits the highest immunogenicity, has the most important role in host interaction, especially cell entry, and is also the main target for neutralizing antibodies [5]. The proteins M, E and HE are only weakly immunogenic and less suitable as targets for antibody diagnosis. Although the N protein is immunodominant, it is not suitable for the specific analysis of the immune response against SARS-CoV-2 viruses due to its high cross-reactivity with antibodies targeting related coronavirius strains [6, 7]. The company Euroimmun AG, Lübeck, Germany offers two ELISA kits using a genetically modified N-protein variant, which enables a more specific detection of antibodies already ten days after infection. However, for highly specific detection of immune response against SARS-CoV-2, typically the S1 subunit of S-protein should be used. Currently, only few vendors offer these specific ELISA formats using the S1 subunit of S-protein for specifically detecting SARS-CoV-2 antibodies [8, 9]. Nevertheless, specific, sensitive and quantitative rapid tests applicable for a decentralized point-of-care (PoC) analysis are currently not available.

Magnetic Immuno-Detection (MInD) could be a powerful tool for PoC assay performance. MInD employs immunofiltration columns (IFCs) containing a porous polyethylene matrix coated with antigens retaining reactive antibodies from applied samples flushed through IFC by gravity flow. Afterward, secondary antibodies specifically binding to the previously enriched antibodies are applied, and subsequently specially functionalized magnetic particles (MNPs) are added, labelling the secondary antibodies. After a washing step to elute unbound MNPs, IFSc are simply inserted into the detection head of a portable magnetic read-out device in which retained MNPs are then detected by means of frequency magnetic mixing detection technology (FMMD), using a low- and a high-frequency magnetic excitation field [10–17]. An alternately oscillating positive and negative magnetic saturation of MNPs with a frequency of 2·*f*_2_ of 122 Hz is obtained when exposing the particle to a low frequency magnetic field of frequency *f*_2_ = 61 Hz with an amplitude of a few millitesla [10]. Afterward, the magnetization state of MNPs is probed by the high frequency magnetic excitation field with *f*_1_ = 49 kHz. An iron oxide dose-dependent signal is obtained when the resulting mixing frequency signal of *f*_1_ + 2·*f*_2_ is detected by a Faraday coil. The innermost coil is composed of two adjacent sections, a so-called detection coil directly encircling the IFC, and a reversely wound empty reference coil, so that the directly induced signal from the excitation coils is cancelled while retaining the nonlinear response signal from the MNPs. A detailed description is found in Ref. [10]. An easy application of this technique is guaranteed due to a direct visualization of the resulting measuring signal at the touchscreen of our portable magnetic read-out device.

We here present a preliminary MInD proof-of-concept study in comparison to a standard laboratory-based ELISA, demonstrating an improved detection of SARS-CoV-2 specific antibodies spiked in human serum samples. We therefore employed a peptide originating from the SARS-CoV-2 Spike protein for IFC coating and antibody enrichment. Our approach might facilitate further optimization to obtain a timely PoC setup for the detection of SARS-CoV-2 specific antibodies in human blood samples.

## 2. Materials and Methods

### 2.1 Material and chemicals

NaCl, KCl, Na_2_HPO_4_ × 12 H_2_0, KH_2_PO_4_, Na_2_(CO_3_), NaHCO_3_, Tris-HCL, MgCl_2_ × 6 H_2_O and Albumin Fraction V (biotin-free) were acquired from Roth, Karlsruhe, Germany. Peroxidase substrate were purchased from Merck KGaA, Darmstadt, Germany.

Coupling buffer was prepared by dissolving 15 mM Na_2_CO_3_ and 35 mM NaHCO_3_ in MilliQ-water, and pH was set to 9.6 with glacial acetic acid. Phosphate buffered saline (PBS) was prepared by dissolving 137 mM NaCl, 2.7 mM KCl, 8.1 mM Na_2_HPO_4_ × 12 H_2_O and 1.5 mM KH_2_PO_4_ in MilliQ-water and setting pH to 7.4 with hydrochloric acid. As washing buffer, PBS with 0.05% (v/v) Tween-20 was used. ELISA and MInD blocking buffer (BP) was prepared by adding 1% (w/v) albumin fraction V (biotin free) in PBS. Alkaline Phosphatase (AP) buffer was prepared by dissolving 100 mM Tris-HCL, 100 mM NaCl and 5 mM MgCl_2_ × 6 H_2_O. Buffer was adjusted to pH 9.6 with HCl. Serum sample from normal healthy human was collected in 2016 and stored at −20°C up to usage.

Immunofiltration columns (IFC) (ABICAP HP columns) were purchased from Senova Gesellschaft für Biowissenschaft und Technik mbH, Weimar, Germany. High-binding 96-well microtiter plates (article number 655061) were purchased form Greiner Bio-One GmbH, Frickenhausen, Germany.

20-aminoacids SARS-CoV-2 spike protein peptide (article number ABIN1382273) derived from the intracellular portion of S2 region of S-protein and corresponding polyclonal rabbit anti-SARS-CoV-2 spike protein peptide specific antibody (article number ABIN1030641) were acquired from antibodies-online GmbH, Aachen, Germany. Biotin-SP (long spacer) AffiniPure Goat Anti-Rabbit IgG (biotinylated GaR secondary antibody), Fc fragment specific (article number 111-065-008) as well as Streptavidin-alkaline phosphatase (streptavidin-AP) (article number 016-050-084) were purchased from Jackson ImmunoResearch Europe Ltd. UK. Streptavidin-functionalized magnetic particles with a hydrodynamic diameter of 70 nm (synomag®-D, article number 104-19-701) were purchased from micromod Partikeltechnologie GmbH, Rostock, Germany.

### 2.2 ELISA Procedure

For ELISA-based SARS-CoV-2 specific antibody detection, all following incubation steps were performed at room temperature for 60 minutes unless stated otherwise. For coating, SARS-CoV-2 spike protein peptide was diluted in coupling buffer to a concentration of 2 μg·ml^−1^ and plated with 100 μl per well onto a high-binding 96-well microtiter plate and incubated. After 3 subsequent washing steps with 200 μl PBS-T per well, each well was blocked with BP and incubated. After washing, 100 μl of SARS-CoV-2 spike protein peptide specific antibody in concentrations ranging from 1.22 ng·ml^−1^ to 5000 ng·ml^−1^ diluted in PBS or in human serum acquired in 2016, respectively, were applied onto the microtiter plate. For blank measurements, PBS or human serum without spiked antibody was employed. After incubation, the plate was washed again. Afterward, 100 μl of biotinylated GaR secondary antibody, diluted 1:20,000 in PBS, was added to each well and incubated. After washing three times with PBS-T, 100 μl of streptavidin-AP (diluted 1:1000 in PBS) was added and incubated for 30 minutes. After washing, absorbance was measured using a Tecan Infinite M200 microplate reader at 405 nm after application of AP-substrate in AP-buffer and 5 minutes of incubation in dark.

### 2.3 Preparation of MInD Immunifiltration Columns

For MInD, IFCs were equilibrated as described by Rettcher *et al.* (2015) [16]. After equilibration, IFCs were coated with 500 μl of 2 μg·ml^−1^ SARS-CoV-2 spike protein peptide, diluted in coupling buffer. After the solution flushed through by gravity flow, a 60 minutes incubation step was performed. Afterward, IFCs were washed by applying 750 μl PBS. Subsequently, remaining binding sites within the polyethylene filter matrix were blocked by applying twice 750 μl BP and an incubation of 20 minutes after each application. After washing of the columns with 750 μl of PBS, IFCs were ready to use for MInD SARS-CoV-2 spike protein peptide specific antibody detection. Pre-coated, ready-to use IFCs could be stored at 4°C for at least several days until they were used for the assay.

### 2.4 MInD SARS-CoV-2 Spike Protein Peptide Specific Antibody Detection

For proof-of-concept MInD-based detection of antibodies against the SARS-CoV-2 Spike protein peptide, dilutions of an antibody with known specificity for this Spike-protein peptide ranging from 1.22 ng·ml^−1^ to 5000 ng·ml^−1^ in PBS or in human serum acquired in 2016, respectively, were prepared and applied onto the ready-to use IFCs. While the sample was flushed through the IFC by gravity flow, SARS-CoV-2 specific antibodies were enriched within the matrix by specific binding to the coated antigen. After 12 minutes of incubation, IFCs were washed with 750 μl PBS and 500 μl of biotinylated GaR (diluted 1:2500 in PBS) were applied onto the columns, binding to retained antibodies. After further incubation of 12 minutes, IFCs were washed again and 500 μl of 60 μg·ml^−1^ 70 nm superparamagnetic streptavidin-functionalized magnetic particles were applied and flushed through the IFCs and incubated for 12 minutes. After a final washing with 750 μl PBS, read-out was done by inserting the columns into the portable magnetic reader, detecting the measuring signal in mV as previously described by Rettcher *et al.* (2015) [16].

### Data Analysis

For ELISA as well as for MInD, data were analyzed using GraphPad Prism 8.0.0, and fitting with a Hill slope was performed. Equations (1) and (2) were used to determine the limit of detection (LOD) or maximum of detection (MOD), respectively, on the signal scale and Equation (3) was used for calculation of those parameters on the concentration scale. SD denotes the standard deviation.

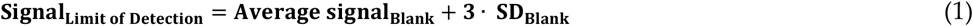

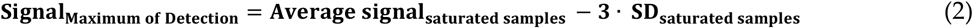

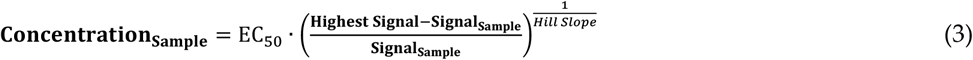

## 3. Results and Discussion

### 3.1. ELISA-based Calibration Experiments of SARS-CoV-2 Specific Antibody Detection

As reference method to our PoC MInD approach, a typical laboratory-based ELISA was performed (Fig 1). After coating of ELISA microtiter plate with SARS-CoV-2 spike protein peptide and blocking with BSA, SARS-CoV-2 spike protein peptide specific antibody was diluted in the range from 1.22 ng·ml^−1^ to 5000 ng·ml^−1^ in PBS-buffer or serum and applied into wells. After addition of biotinylated GaR and subsequent labelling with streptavidin-AP, the ELISA plate was read out at 405 nm and obtained measuring values were used to generate calibration curves for SARS-CoV-2 specific antibody concentrations in PBS (Fig 1, black curve) and in human serum samples (Fig 1, red curve). Blank values determined in PBS and serum were 0.085 AU ± 0.005 AU and 0.083 AU ± 0.001 AU, respectively, and were subtracted from sample values. Both curves recorded with antigen diluted in PBS and human blood serum were almost identical. Based on a Hill fit, calibration measurements in PBS with a LOD 3.4 ng·ml^−1^ and or LOD of 4.0 ng·ml^−1^ in human serum were obtained (Fig 1). Both calibration measurements show saturation of measuring signals at concentrations with 625 ng·ml^−1^ up to 5000 ng·ml^−1^. For calculation of maximum of detection, the average measuring signal of samples with concentrations ranging from 625 ng·ml^−1^ up to 5000 ng·ml^−1^ was calculated and threefold SD was subtracted. Using equation 3, a MOD of 477 ng·ml^−1^ in PBS and of 312 ng·ml^−1^ in serum, respectively, could be determined. Following these calibration measurements, it can be concluded, that with our laboratory-based ELISA, SARS-CoV-2 specific antibodies can be detected in range of 3.4 ng·ml^−1^ up to 477 ng·ml^−1^ in PBS-buffer or from 4.0 ng·ml^−1^ up to 312 ng·ml^−1^ in human serum samples, respectively. Typical IgG antibody concentrations in human serum are approximately 10 mg·ml^−1^, and are significantly increasing after antigenic stimulation of immune system [18]. For the whole assay, excluding coating and blocking but including application of sample, secondary antibody, streptavidin-AP and substrate incubation, a procedure time of 161 minutes was needed. This time is comparable to that of commercially available ELISA test kits (e.g. Euroimmun 140 min). Comparing the sensitivity of commercially available ELISA test kits with our standard ELISA will be possible when samples of COVID-19 patients are evaluated. A highly sensitive assay could detect antibodies at an early stage of infection, whereas commercially test kits can detect IgG antibodies in patient samples against S1 subunit of SARS CoV-2 S-protein in 75% of samples 10 to 20 days after infection (Euroimmun).

**Fig 1.**
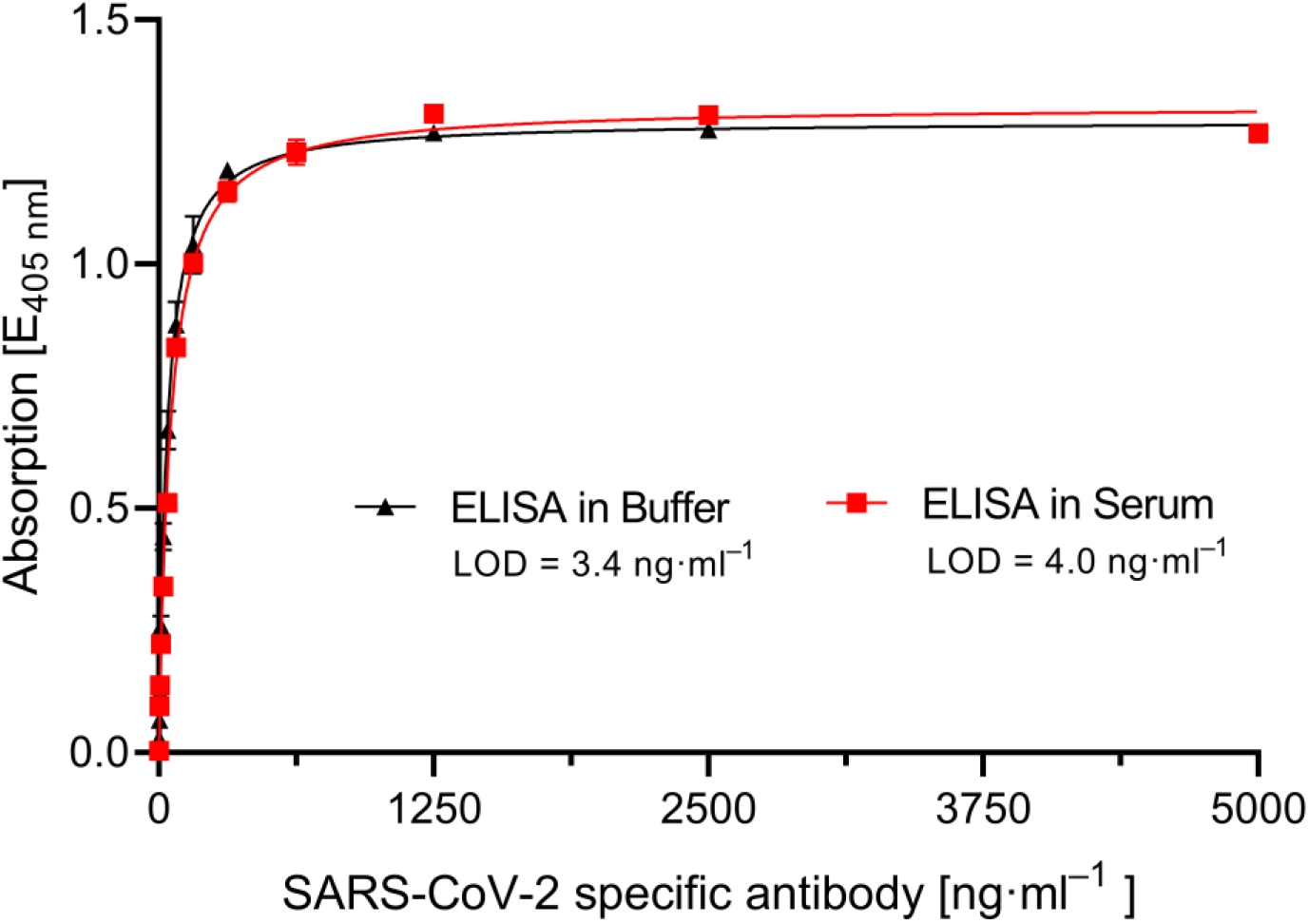
**ELISA-based detection of SARS-CoV-2 spike protein specific antibody** in PBS-buffer (black curve) or spiked in human serum (red curve). Antibody was diluted to concentrations ranging from 1.22 ng·ml^−1^ up to 5000 ng·ml^−1^ in each matrix and applied onto 2 ng·ml^−1^ SARS-CoV-2 spike protein peptide coated and BSA blocked microtiter plates. After addition of biotinylated secondary antibody, streptavidin-AP was applied. Limit of detection (LOD) was calculated using non-linear Hill fit (R2=0.997 for PBS-buffer and 0.996 in serum). Assay time of ELISA was 161 minutes. Each data point represents mean ± SD (n = 4 for PBS-buffer and n = 3 for serum).

### 3.1. MInD-based Calibration Experiments for SARS-CoV-2 Specific Antibody Detection

Same calibration measurements employing dilutions of SARS-CoV-2 specific antibody were done with our PoC MInD-based setup (Fig 2 and 3). Comparable to laboratory-based ELISA, the same dilutions of SARS-CoV-2 spike protein peptide specific antibody in PBS-buffer (Fig 3, black curve) or human serum (Fig 3, red curve) were applied after coating and blocking of IFC with SARS-CoV-2 spike protein peptide and BSA. After application of antibody dilutions, a 5-times shorter incubation time of just 12 minutes compared to ELISA was performed. The reduction of incubation time could be achieved due to a more efficient antibody enrichment. The surface for target binding within the ABICAP immunofiltration column matrix is approximately 40-fold larger compared to the surface of an ELISA well. Furthermore 500 μl of sample was applied and flushed through IFC, compared to just 100 μl of sample added to an ELISA microtiter plate well. Afterward, biotinylated GaR was applied onto columns and magnetically labelled with 70 nm streptavidin-functionalized magnetic particles. Again, incubation time of only 12 minutes each could be used due to the increased binding surface. Finally, the IFCs were inserted into our portable magnetic read-out device which can be operated using either an external power adapter or a portable battery, allowing a PoC diagnostic assay procedure. A schematic overview of assay procedure is shown in Fig 2.

**Fig 2.**
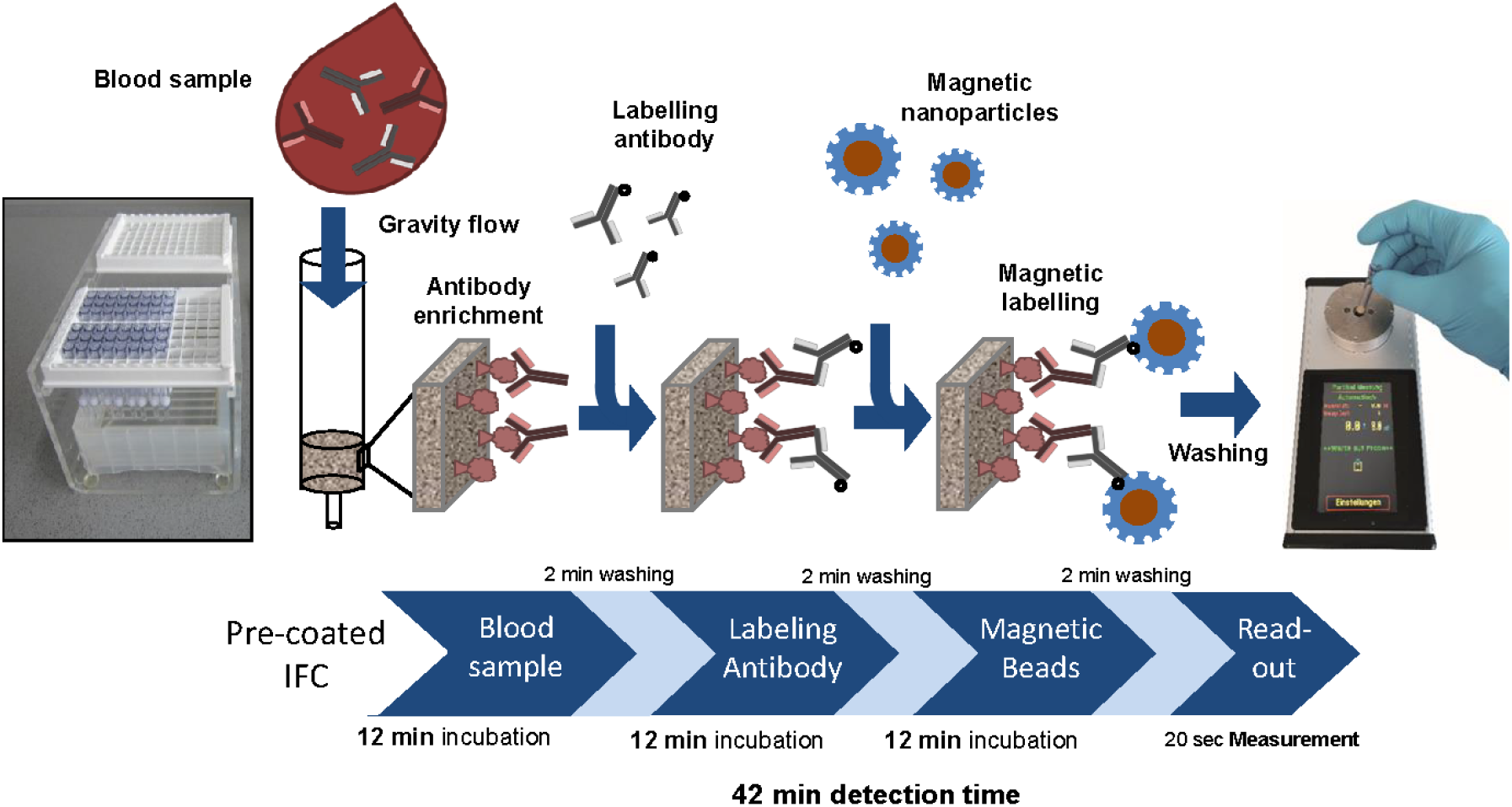
Proof-of-concept MInD assay setup using IFC coated with SARS-CoV-2 antigen. Assay steps and assay time are indicated. IFCs were coated with commercial SARS-CoV-2 S-protein peptide and blocked with BSA. Corresponding antibody was diluted either in PBS or spiked in human serum and applied to IFCs. Biotinylated secondary antibodies were added, followed by application of streptavidin-functionalized MNP. Finally, IFCs were inserted into the portable magnetic read-out device. Measuring signal can be correlated to the amount of antibody in the sample and antibody titer can be determined. Assay time of this preliminary MInD setup was 42 min which is approximately four times faster than ELISA (161 min).

**Fig 3.**
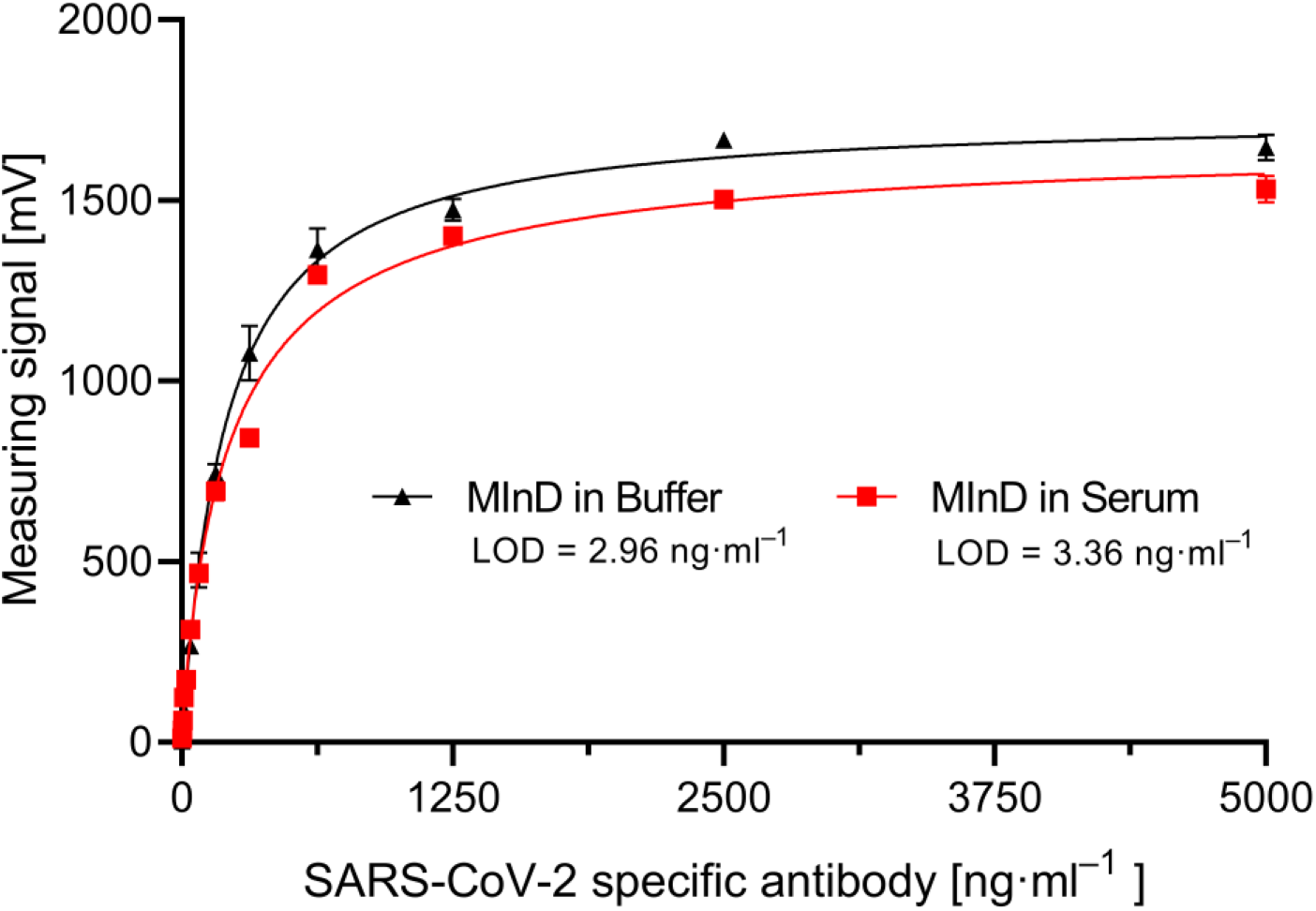
Proof-of-concept MInD-based SARS-CoV-2 specific antibody detection. IFCs were coated with commercial 2 μg·ml^−1^ SARS-CoV-2 spike protein peptide and blocked with BSA. A corresponding antibody was diluted either in PBS (black curves) or spiked in human serum (red curves) and applied to IFCs. Biotinylated secondary antibodies were added, followed by application of streptavidin-functionalized magnetic particles. Assay time of this preliminary MInD setup was 42 minutes (without column preparation). Limit of detection (LOD) was determined by help of non-linear hill fit. Each data point represents mean ± SD (n = 2).

As shown in Fig 3, also with MInD-based approach SARS-CoV-2 specific antibody detection calibration curves could be recorded using PBS-buffer or human serum, respectively. Particularly the comparable calibration measurements in PBS or human serum demonstrate the negligible matrix effect of human serum when using MInD. Compared to standard ELISA, a saturation of measurement signal was observed only at samples with higher concentrations of 2500 ng·ml^−1^ and 5000 ng·ml^−1^. Based on this, the average of these two samples was used for calculation of the MOD (Equation 2 and 3). By analyzing the range of detection, it can be seen that SARS-CoV-2 spike protein peptide specific antibody can be detected in range of 2.95 ng·ml^−1^ up to 2040 ng·ml^−1^ in PBS and from 3.36 ng·ml^−1^ up to 1810 ng·ml^−1^, demonstrating an at least 5-fold broader range of quantification in both PBS and human serum with higher sensitivities compared to ELISA and perfectly matching nonlinear fit (R^2^ =0.997 in PBS and R^2^ =0.993 in human serum). This significantly increased dynamic range compared to ELISA demonstrates one major advantage enabling improved quantitative measurements.

Furthermore, an approximately 4-fold reduction of assay time with PoC MInD approach compared to ELISA demonstrates the high potential for fast assay procedure. In total, 42 minutes procedure time was needed, resulting in a broader detection range in combination with lower detection limits in both PBS and human serum. A further reduction of assay time could be achieved by checking the required amount of washing steps and analyzing the achieved sensitivity when reducing the incubation time to 5 minutes or less, as described by Rettcher *et al.* (2015) [16]. Additionally, Rettcher and colleagues demonstrated a reduction of assay time by pre-functionalizing the MNPs with antigen-specific antibodies [16]. Here, MNPs can be pre-functionalized with secondary antibody, which would result, in combination with previously mentioned optimization steps, in less assay steps with a comparable assay time as PoC lateral flow assays (less than 20 minutes).

In this proof-of-concept experiment, a commercially available SARS-CoV-2 spike protein peptide with corresponding antibody was used. If using this peptide for testing of patient samples, there might be a high risk for false negative assay results, since patients may not have developed antibodies against this peptide. Most preferably, other antigen variants derived from the highly immunogenic S1-subunit of S-protein or a mixture of antigens of SARS-CoV-2 proteins should be used for highly effective and specific enrichment of SARS-CoV-2 targeting antibodies. Especially for demonstration of MInD specificity, control antigens derived from common cold human coronaviruses (hCoV) as hCoV 229E, hCoV NL63, hCoV OC43, hCoV HKU1 or SARS-CoV and MERS-CoV should be tested, confirming the enrichment of only SARS-CoV-2 specific antibodies. Additionally, a multiplex detection of different MNPs, as demonstrated by Achtsnicht *et al.* (2019), could be implemented for detection of multiple antibody subclasses in one assay [19]. By coupling different secondary antibodies to MNPs, each type will label a specific antibody isotype, as e.g. IgA and IgG, a course of infection could be visualized as well as the analysis of seroconversion would be enabled. A further optimization of the magnetic read-out device towards a medical-diagnostic device would then fulfill all requirements for use in medical field. To ensure correct sample allocation, the magnetic reader can be equipped with a barcode scanner, and IFCs could be labelled with a patient-specific barcode. In combination with multiplex detection of several antibody isotypes, the MInD approach would thus be ideally suited for the use in doctors’ surgeries since our MInD assay has less than 10% of the cost for typical ELISA equipment. Especially due to the possibility of performing the PoC assay procedure with a single pipette, and the portable magnetic read-out device which can be battery-operated, the MInD assay can also be used by service providers in the medical field and for testing in elderly peoples’ and nursing homes or at airports, quickly identifying persons with existing immunity. As soon as vaccination against SARS-CoV-2 is available, our approach could be employed to monitor vaccination success and longevity of immunity by determining antibody titers.

## 4. Conclusions

We demonstrated for the first time a proof-of-concept MInD-based approach for rapid and highly sensitive SARS-CoV-2 S-protein peptide specific antibody detection in spiked human serum. MInD calibration experiments with a five-fold higher range of detection in combination with higher sensitivity and a four-fold shorter assay time in comparison to a standard ELISA demonstrate the high potential of MInD-based PoC SARS-CoV-2 specific antibody detection in serological samples. By using appropriate SARS-CoV-2 antigens and a multiplex approach for simultaneous detection of e.g. IgA and IgG antibodies reactive against SARS-CoV-2, the state of infection as well as a seroconversion could be analyzed. Especially due to assay speed, low-cost and portable equipment, we conclude that the MInD-based assay would be ideally suited for PoC testing, identifying persons with existing immunity. Additionally, our MInD approach could be employed for subsequent analysis of efficiency and for monitoring the longevity of vaccination by determining antibody titers.

## Author Contributions

Conceptualization, F.S. and J.P.; methodology, J.P. and N.V.; validation, J.P.; formal analysis, J.P.; Resources, F.S., H.S. and H.J.K; investigation, F.S. and J.P.; data curation, J.P.; writing—original draft preparation, J.P.‮; writing—review and editing, F.S., H.J.K, H.S. and N.V.; visualization, J.P.; supervision, F.S.; project administration, F.S.

## Acknowledgments

The authors would like to thank Max Schubert for his helpful advices and support given in discussions.

